# Can leafhoppers help us trace the impact of climate change on agriculture?

**DOI:** 10.1101/2023.06.13.544773

**Authors:** Nicolas Plante, Jeanne Durivage, Anne-Sophie Brochu, Tim Dumonceaux, Dagoberto Torres, Brian Bahder, Joel Kits, Antoine Dionne, Jean-Philippe Légaré, Stéphanie Tellier, Frédéric Mcune, Charles Goulet, Valérie Fournier, Edel Pérez-López

## Abstract

**SUMMARY:** Climate change is reshaping agriculture and insect biodiversity worldwide. With rising temperatures, insect species with narrow thermal margins are expected to be pushed beyond their thermal limits, and losses related to herbivory and diseases transmitted by them will be experienced in new regions. Several previous studies have investigated this phenomenon in tropical and temperate regions, locally and globally; however, here, it is proposed that climate change’s impact on agriculture can be traced through the study of Nearctic migratory insects, specifically leafhoppers. To test this hypothesis, leafhoppers in strawberry fields located in the province of Québec, eastern Canada, were evaluated. The strawberry-leafhopper pathosystem offers a unique opportunity because leafhoppers can transmit, among other diseases, strawberry green petal disease (SbGP), which is associated with pathogenic phytoplasmas. Here, we found that in the last ten years, the number of leafhoppers has been increasing in correspondence with the number of SbGP cases detected in eastern Canada, reporting for the first time ten species new to eastern Canada and two to the country, although the leafhopper diversity has been seriously affected. Our model using more than 34 000 leafhoppers showed that their abundance is influenced by temperature, a factor that we found also influences the microbiome associated with *Macrosteles quadrilineatus*, which was one of the most abundant leafhoppers we observed. One of our most striking findings is that none of the insecticides used by strawberry growers can control leafhopper incidence, which could be linked to microbiome changes induced by changing temperatures. We suggest that Nearctic leafhoppers can be used as sentinels to trace the multilayered effects of climate change in agriculture.

**GRAPHICAL ABSTRACT:** 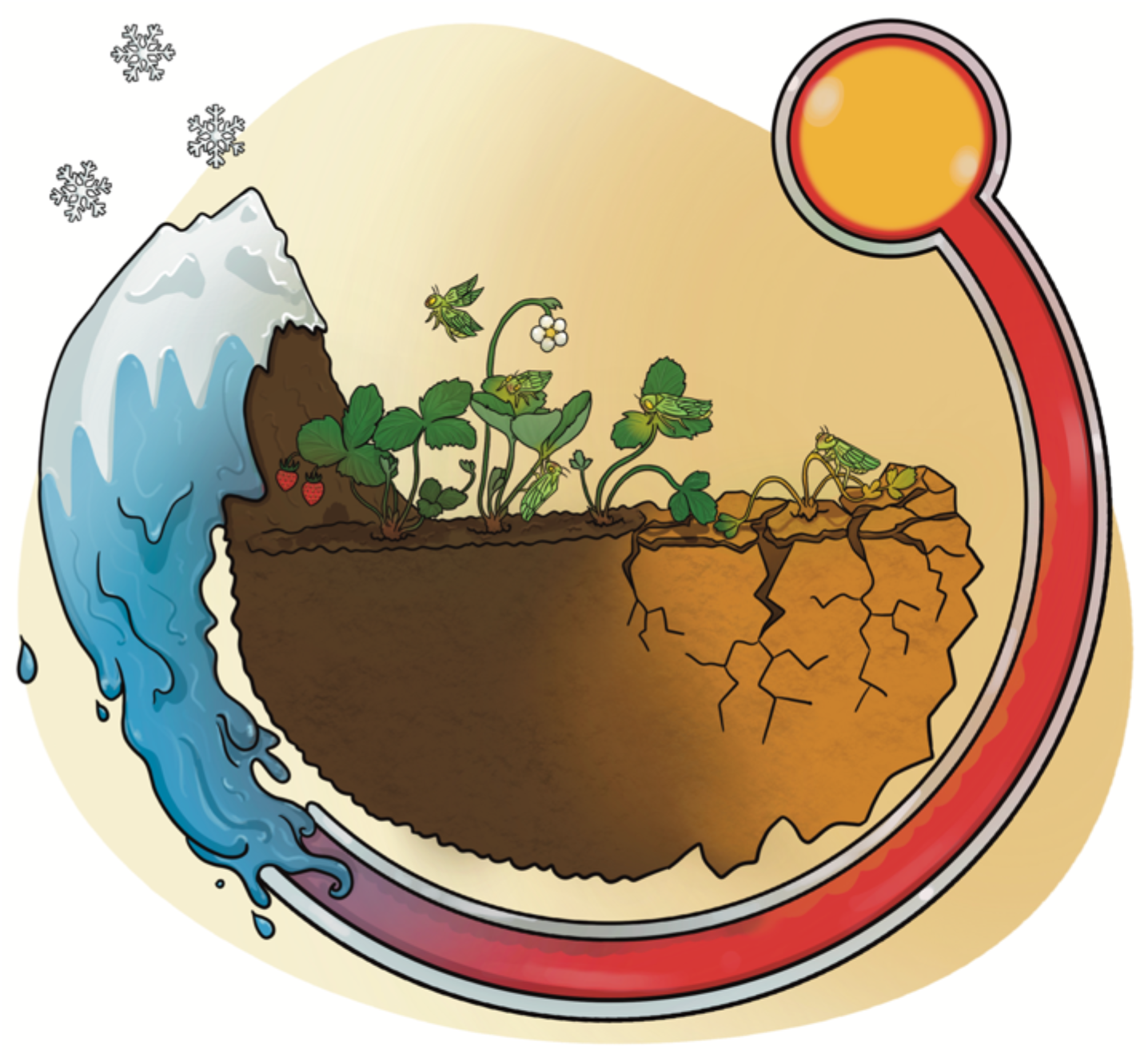

**IN BRIEF:** The current climate crisis is reshaping insect biodiversity and abundance, but little is known about the direct effect of this phenomenon on agriculture. In this study, we explored leafhoppers, a group of agriculturally important insect pests and disease vectors, as sentinels of the effect of climate change on agriculture. Our findings indicate that this group of insects can help us to understand the effect of the current climate crisis on insect invasions, diversity, abundance, disease dynamics and insecticide resistance and to take quick action to ensure food security while achieving more sustainable agriculture.

**HIGHLIGHTS:**

- Migratory leafhoppers benefit from temperature increases
- Leafhopper-transmitted diseases have increased in the last decade
- New non-migratory leafhoppers can be found now in Nearctic regions
- Leafhopper insecticide resistance could be linked to the insect microbiome

## 1. INTRODUCTION

Climate change is a major threat to agriculture (Raderschall et al., 2020). In the coming years, a warmer climate is expected, which will have a serious impact on insect pest distribution and disease incidence, posing a risk to food security (Ristaino et al., 2021; Outhwaite et al., 2022). The impact of climate change on insect biodiversity is complex and variable depending on the taxonomic group (Jactel et al., 2019; Lehmann et al., 2020; Ristaino et al., 2021; Outhwaite et al., 2022). Increases in temperatures promote biological invasions by expanding the thermal limits of various insect populations (Harvey et al., 2020; Rossi & Rasplus, 2023), a phenomenon particularly relevant in areas that have been sheltered from pests principally due to climatic differences, as is the case in the northern territories of the Nearctic region (Colinet et al., 2015). Higher temperatures also increase the number of generations of insects per year (Deutsch et al., 2008; Harvey et al., 2020) and insect metabolic rates, accelerating phytophagy (Dillon et al., 2010; Deutsch et al., 2018). However, things become more complex when we analyze the effects of climate change on plant-vector-pathogen interactions. At the level of the pathogen, there is evidence that some groups of insect-transmitted bacteria, such as phytoplasmas, can multiply faster in the plant host at higher temperatures, increasing the chances of transmission by the vector during sap feeding (Maggi et al., 2014; Bahar et al., 2018; Sabato et al., 2020). Increased phytophagy, a higher number of insect generations per year, and higher availability of pathogens in the host will directly contribute to a higher dispersion of pathogens and diseases. These perturbations will affect plant productivity and might require extreme intervention and the use of higher amounts of pesticides (Garret et al., 2006; Ma et al., 2021). Unfortunately, we know today that climate warming will also contribute to insecticide resistance through different mechanisms (Almeida et al., 2017; Ma et al., 2021; Gupta et al., 2022). By expanding their overwintering range, insect pests can develop insecticide resistance more quickly through frequent gene exchange in non-overwintering sites (Ma et al., 2021), while warmer temperatures can also promote shifts in insect microbiomes that contribute to better adaptation to new conditions and insecticides (Wang et al., 2022; Gupta et al., 2022; Zhang et al., 2023). Although all these effects of climate warming have been studied using a diverse array of insects as models, the effects on plant-vector-pathogen interactions are less understood, and no suitable pathosystem has been identified that could allow us to see the expected multilayered effects.

Leafhoppers (Cicadellidae) are a group of phloem-, xylem- and mesophyll-feeding insects that includes more than 20,000 described species and is the second-largest hemipteran family in the world (Stiller 2009). These insects are also well-known vectors of viral and bacterial diseases affecting a wide range of economically and ecologically important crops (Weintraub & Beanland, 2006; Brochu et al., 2023). Among those pathogens transmitted by leafhoppers, phytoplasmas, a group of unculturable, cell wall-less bacteria that are members of the class Mollicutes, are probably the most studied (Bertaccini, 2022). Leafhoppers within ten subfamilies and more than 70 species have been identified as vectors of phytoplasma diseases worldwide (Weintraub & Beanland, 2006; Silva-Castaño et al., 2023), but the understanding of these tritrophic relationships is in its infancy. A group of leafhoppers that has been relatively well studied are Nearctic leafhoppers, referring to those distributed from Mexico to the Arctic (Omar 1949). This group comprises close to 3000 species that are divided into almost 20 subfamilies and includes several well-known phytoplasma vectors, such as *Macrosteles quadrilineatus* (Forbes, 1885), the aster yellows leafhopper and *Empoasca fabae* (Harris), the potato leafhopper (Ishii et al., 2013; Baker et al., 2015). Some Nearctic leafhoppers have a very well-delimited niche, which is influenced by temperature, and in some cases, such as the genus *Dalbulus*, by a strong host specificity (Bernal et al., 2019). However, most of the species in this geographic group are oligophagous, with many polyphagous species, and their distribution is mainly controlled by food availability and thermal limits.

Canada contains only 5% of the reported global leafhopper diversity and 35% of Nearctic leafhoppers (Maw et al., 2000) due to very cold winters and limited food availability. Climate change is expected to impact the numbers and diversity of leafhoppers that can live and reproduce in the northern reaches of the Neartic range (Firlej et al., 2019). The annual average temperature in Canada has increased at approximately twice the global mean rate, and direct effects of warming, such as extreme heat, less extreme cold, longer growing seasons, shorter snow and ice cover seasons, and earlier spring peaks, are increasingly common (Bush & Lemmen, 2019). These changes in temperatures, based on previous studies, will expand the thermal limits of many Nearctic leafhopper species, but little research has been done on the subject. How can these effects be quantified in agriculture and plant-vector-pathogen interactions? In a recent study, it was determined that, only in the province of Québec, the number of hosts affected by phytoplasma has quintupled in the last decade (Brochu et al., 2023), evidence that the number of vectors might have increased, that new leafhopper species could be contributing to the spread of the pathogens, and/or that the management strategies used to control the vectors are not successful. Here, to identify what scenario(s) could be taking place, we studied the strawberry-leafhopper-strawberry green petal phytoplasma pathosystem (Brochu et al., 2021). After two growing seasons, we found that this pathosystem is strongly influenced by changes in temperatures and that what we thought was a local study, could have a great impact on the way we trace the impacts of climate change on agriculture and how we will manage the imminent biological invasions already happening in places such as Canada.

## 2. MATERIALS AND METHODS

All data analyses and graphs were made using the R language for statistical computing (R Core Team, 2021). Below, we specify the package used for each analysis (Hothorn et al., 2008; Bates et al., 2015; Oksanen and al., 2022; Pinheiro & Bates, 2023; Lenth, 2023; Mazerolle, 2023).

### 2.1. Experimental setting

This study was conducted in seven strawberry fields distributed in four geographic regions in the province of Québec, Canada: Capitale-Nationale, Chaudière-Appalaches, Mauricie and Montérégie, during the growing seasons from 2020 to 2022 (Supplementary Table S1). Strawberry fields were composed of six strawberry varieties in different combinations (Supplementary Table S1), and leafhopper capture was performed with yellow sticky traps spatially distributed to cover the field. Each trap was spaced one meter apart. Yellow traps were changed weekly from the third week of May to the third week of September of each growing season. During the first growing season, we placed 90 traps per week, while in the second growing season, to accommodate the dimensions of the fields, we placed 75 traps per week.

### 2.2. Leafhopper identification

Sticky traps were collected from the field, and leafhoppers were removed from the traps and stored at 4°C in Eppendorf tubes with 70% ethanol until further identification. Leafhopper identification was performed based on morphological comparisons of male genitalia using taxonomic keys of each genus. Abdomens were removed from all male specimens, and the genitals were directly dissected or placed into a 10% KOH solution of tissue lysis buffer for 24 hours at 56°C to remove unsclerotized material and fat. Female specimens were mostly identified at the genus level, except for some easily identifiable species and those that did not significantly differ from males. Identified specimens were compared to reference material at the Canadian National Collection of Insects, Arachnids and Nematodes (Agriculture and Agri-Food Canada, Ottawa, Ontario, Canada) and a subset confirmed by the National Identification Service. As a confirmatory analysis, specimens from eighteen genera were randomly selected and identified molecularly using the primer pair LCO1490/HCO2198 amplifying the COI gene as previously described (Leray et al., 2013). Pictures of the body and genitals of the leafhopper were taken using a Leica M205 C stereomicroscope. The COI sequences generated in this study were deposited in GenBank.

### 2.3. Leafhopper abundance, diversity, and population distribution

To evaluate leafhopper diversity, alpha diversity indices were first used to determine differences in diversity between years and among geographic regions. To compare the sites with each other, we normalized the number of leafhoppers captured each year according to the number of traps per site and the time that each trap remained in each field. We determined the Shannon (H’) diversity index, Simpson (D) dominance index and species richness for each region and year of sampling with the R vegan package (Oksanen et al., 2022). Then, we investigated if there was a significant difference in diversity between years, among regions, and finally, if there was an interaction between regions and years using repeated measures ANOVA with sampling sites as a random source.

To investigate whether temperature could be used to predict leafhopper abundance, we constructed a negative binomial mixed model with historical temperature and rain data collected from Environment Canada’s historical database (https://climat.meteo.gc.ca/historical_data/search_historic_data_e.html). We selected the weather stations that were closest to the sampling locations (Supplementary Table S2). The bioclimatic model considered the site as a random effect, the time of year as a fixed effect, and a temperature-related index. The model considered the number of traps and the time length of exposure of the traps as an offset for each entry. Five temperature-related indexes were tested in different models: (*i*) the mean temperature over each period of leafhopper capture and the contribution of the period to the annual total degree-days (*ii*) base 2 °C, (*iii*) base 5 °C, (*iv*) base 8 °C and (*v*) base 10 °C (Trudgill et al., 2005). Different polynomial models were made, with all possible combinations of first, second and third degrees for the temperature and the time factors, independently, and for each temperature-related index. The choice of the best model was based on the Bayesian information criterion (BIC). This procedure was then repeated with *Macrosteles quadrilineatus* and *Empoasca fabae* abundance data. Precipitation was added to the best model with all leafhopper species. It was tested as a first, second and third polynomial degree factor. These models were compared with the model without rain based on their BICs to determine if rainfall is a relevant factor.

### 2.4 Phytoplasma characterization

To investigate the phytoplasma associated with strawberry green petal disease (SbGP) in Eastern Canada, twelve plants showing typical symptoms of the disease between the two growing seasons (Supplementary Table S3) (Brochu et al., 2021) and one asymptomatic plant were collected from three strawberry fields located at Capitale-Nationale, Québec, Canada. Genomic DNA was extracted from the leaf midrib using a CTAB-based method (Murray & Thompson, 1980) and used as a template for phytoplasma characterization by multilocus sequence typing as previously described (Pusz-Bochenska et al., 2022). The sequences of the 16S, *cpn60*UT, *tuf*, *secY*, *secA*, *nusA*, and *rp* phytoplasma taxonomic markers generated using this method from the twelve symptomatic samples were used to determine phylogenetic relationships with MEGA v. X (Kumar et al., 2018) and the neighbour-joining method with 1000 bootstraps.

In parallel, genomic DNA from three symptomatic plants was used as a template for direct PCR. with phytoplasma universal 16S primers R16F2n/R16R2 (Gundersen & Lee, 1996) and *cpn60UT* with primers H279p/H280p (Dumonceaux et al., 2014) using the high-fidelity polymerase Phusion Plus (Thermo Scientific). Amplicons were obtained from the three symptomatic samples (named SbG, SbG and SbG), while no amplification was obtained using genomic DNA from asymptomatic plants. The bands generated through PCR from all samples were cloned into the vector pGEM-T Easy (Promega), and the recombinant plasmid was transformed into chemically competent *Escherichia coli* TOP10 (Life Technologies). Two to three clones from SbG, SbG and SbG were sequenced using plasmid-targeted M13 sequencing primers (forward and reverse). The sequences generated from the samples were aligned and analyzed using the *i*Phyclassifier (Zhao et al., 2013) and CpnClassiPhyR (Muirhead et al., 2019) online tools to determine ‘*Candidatus* Phytoplasma’ species, as well as group and subgroup classification of the SbGP strain detected. These sequences were also used to construct a neighbour-joining phylogenetic tree as described above. All the phytoplasma-derived sequences generated in this study were deposited in GenBank. Additionally, to investigate the distribution and incidence of the disease, we collected historically reported data from the Québec government (Ministry of Agriculture, Fisheries and Food of Québec) tracing the disease in the last decade (Tellier & Lacroix, 2012) (Supplementary Table S4).

### 2.5. Phytoplasma vector identification

To determine which leafhopper could be an SbGP phytoplasma vector, we followed three strategies:

#### 2.5.1. Phytoplasma amplification from captured leafhoppers

For this, we extracted genomic DNA from 10% of the leafhoppers collected during the first growing season. The 2926 leafhoppers were grouped into samples of one to 10 specimens (Supplementary Table S5), and DNA was extracted using a CTAB-based method (Murray & Thompson, 1980) with a buffer concentration of 2% CTAB, 100 mM Tris-hydrochloric acid (HCl), pH 8.0, 20 mM ethylenediaminetetraacetic acid (EDTA), pH 8.0 and 1.4 M NaCl. Phytoplasma detection was performed using DNA as a template in a direct PCR with phytoplasma universal 16S primers R16F2n/R16R2 (Gundersen & Lee, 1996). PCR products were analyzed and visualized by electrophoresis in a 1% agarose gel with pGEM-T Easy containing the SbGP-derived 16Sr operon as the positive control and double distilled water as a negative control.

#### 2.5.2. Incubation of strawberry plants with leafhoppers

From June to August, the peak of leafhopper abundance of each growing season, we captured leafhoppers using an entomological aspirator and placed them in cages with disease-free strawberry plants obtained from seeds. Leafhoppers were incubated for seven to 14 days with the plants, and after the insects were captured back to perform DNA extraction, they were separated by genus or species in samples consisting of one to five specimens. This was performed 22 times between both growing seasons, and a total of 278 leafhoppers within eight genera were analyzed (Supplementary Table S6). In parallel, genomic DNA was also extracted from the leaf midrib, and leafhopper DNA was used as the template for phytoplasma detection by 16S PCR amplification as described above.

#### 2.5.3. <u>Macrosteles quadrilineatus</u> microbiome analysis

To study the microbiome composition of *Macrosteles quadrilineatus* (Forbes, 1885), the aster leafhopper and well-known phytoplasma vector, 300 specimens were analyzed from each growing season. For each season, we analyzed 150 leafhoppers collected in June and 150 collected in August. These were collected in three geographic regions: (*i*) Montéregie, (*ii*) Mauricie, and (*iii*) Capitale Nationale, with 50 leafhoppers per region separated into five samples consisting of ten leafhoppers each (Supplementary Table S7).

Using the aster leafhopper genomic DNA as a template, the V4-V5 region of the 16S rRNA universal target was amplified using 30 cycles of 94°C for 45 s, 50°C for 30 s, and 72°C for 90 s. The primers used were 515F (Parada et al., 2016) and 926R (Quince et al., 2011). Amplifications used 1X PCR Buffer (Thermo Fisher Scientific, MA, USA), 2.5 mM MgCl2, 0.5 mM dNTPs, 0.4 µM of each primer and 1 U of Platinum Taq polymerase (Thermo Fisher Scientific, MA, USA) in a total volume of 25 µl. Amplification negative controls containing only buffers and water were processed and sequenced along with experimental samples. Indexed 16S amplicons were pooled into a single sample of 4 nM total and sequenced using 300 PE cycles of Illumina MiSeq Chemistry (Illumina, San Diego, CA, USA).

Cutadapt (v.2.8) (Martin, 2011) was used to remove amplification primer sequences and trim bases with a quality score < Q30. Sequencing reads were merged using FLASH2 (v.2.2) (Magoč & Salzberg, 2011). Merged reads were used to define amplicon sequence variants (ASVs) using the q2-DADA2 (Callahan et al., 2016) plugin for QIIME2 (v.2020.2) (Bolyen et al., 2019). Taxonomic classification of the ASVs used Naïve Bayes Classification with the q2-classifier in QIIME2 and the silva-138-99 (Quast et al., 2012) reference database. ASVs that were classified as mitochondrial or chloroplast sequences were removed.

Microbial community diversity statistics, including Bray-Curtis dissimilarity and Shannon diversity (H’), were calculated after rarefaction to the smallest library size (6493 reads for the 2021 data; 2622 reads for the 2022 data) with the phyloseq (McMurdie et al., 2013) and vegan (Oksanen et al., 2007) packages in R. Differences in alpha and beta diversity were tested for significance using the Kruskal-Wallis test (Benjamini-Hochberg false discovery rate correction) and permutational multivariate analysis of variance (PERMANOVA), respectively. The microbiome data generated in this study were deposited in GenBank.

### 2.6. Efficiency of leafhopper management

To investigate the efficiency of insecticides in controlling leafhopper populations, we compared the effects of different insecticides used by strawberry growers and registered as effective against leafhoppers in the reduction of these insect pests (Supplementary Table S8). The abundance of leafhoppers before and after each insecticide treatment was statistically compared using repeated measures ANOVA to determine whether insecticides were significantly effective for controlling leafhoppers. We analyzed the insecticide’s effect on all leafhopper species captured and on *Empoasca fabae* (Harris) and *M. quadrilineatus* independently.

## 3. RESULTS

### 3.1. Leafhopper abundance and diversity

In total, we collected 33 007 leafhoppers in the strawberry fields sampled during this study (Fig. 1A, Supplementary Table S9). In both seasons, leafhopper populations began to increase from mid-June and reached a maximum peak around mid-July (Fig. 1B). In each region, *Macrosteles* spp., *Empoasca* spp., and *Hebata* spp. were consistently among the most abundant genera (Fig. 1C-F). The potato leafhopper and the aster leafhopper alone represented 88.4% of all specimens captured during the first growing season and 77.4% of all captures during the second (Supplementary Table S9).

**Figure 1.**
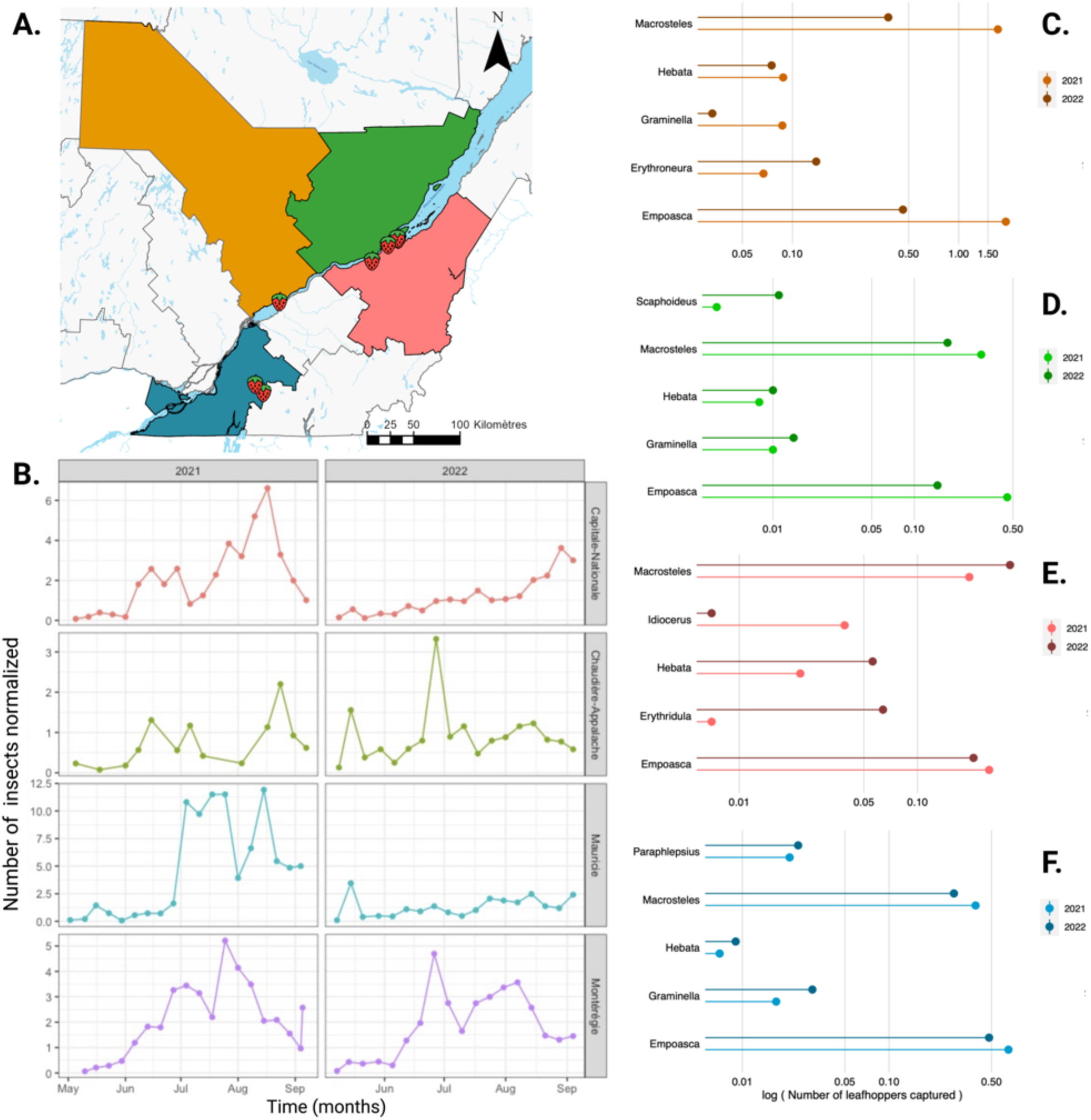
Leafhopper abundance reported in this study. **A**, geographical location of the strawberry fields screened in this study. Colours signal the four geographic regions where the fields are located, with blue for Montérégie, orange for Mauricie, red for Chaudière-Appalache, and green for Capitale-Nationale. **B**, Leafhopper abundance distribution from May to September during both growing seasons, hereafter called 2021 and 2022. **C**-**F**, the five most abundant leafhopper genres captured in each region with **C** for Montéregie, **D** for Chaudière-Appalache, **E** for Capitale-Nationale, and **F** for Mauricie. The number of insects was weighted based on the number of traps and the number of days that each trap remained in the field in all analyses.

After taxonomic identification, 61 genera and 118 species within seven subfamilies were found among the leafhoppers captured in strawberry fields in this study (Fig. 2A-R, Supplementary Table S9). In all cases, the molecular identification of the 18 randomly selected genera was consistent with the morphology-based identification, except for *Xestocephalus fulvocapitatus* (Evans, 1954), which is the first CO1 sequence registered in GenBank for this species (Fig. 2S). Among all the species captured, 11 species were identified as new to the province of Québec, ten were reported the furthest east ever observed in Canada, and one was reported for the first time in Canada (Fig. 2T, Supplementary Table S9). The new reports for Québec and eastern Canada are *Erythridula plena* (Beamer), *Graminella pallidula* (Osborn), *Macropsis fuscula* (Fieber), *Erythroneura vitifex* (Fitch), *Erythroneura elegans, Erythroneura rubrella* (McAtee), *Hebata vergena* and *Hebata ditata* (DeLong & Caldwell), while *Erythridula wysongi* (Ross & DeLong) and *Erythroneura bakeri* (Dmitriev & Dietrich) are new reports for Canada. In addition, we found that the diversity was significantly higher during the second growing season based on the Shannon (*p* = 0.0151) and Simpson (*p* = 0.0417) diversity indexes (Fig. 2U, Supplementary Table S10). Curiously, although the diversity followed a similar trend among the regions using both diversity indexes, only when the diversity was estimated with the Shannon index did we observe significant differences (*p* = 0.0275), with Capitale-National showing the highest diversity among the regions (Fig. 2U, Supplementary Table S10). Finally, no significant effect was detected on the interaction between regions and years or on species richness (Supplementary Fig. S1 and Table S10).

**Figure 2.**
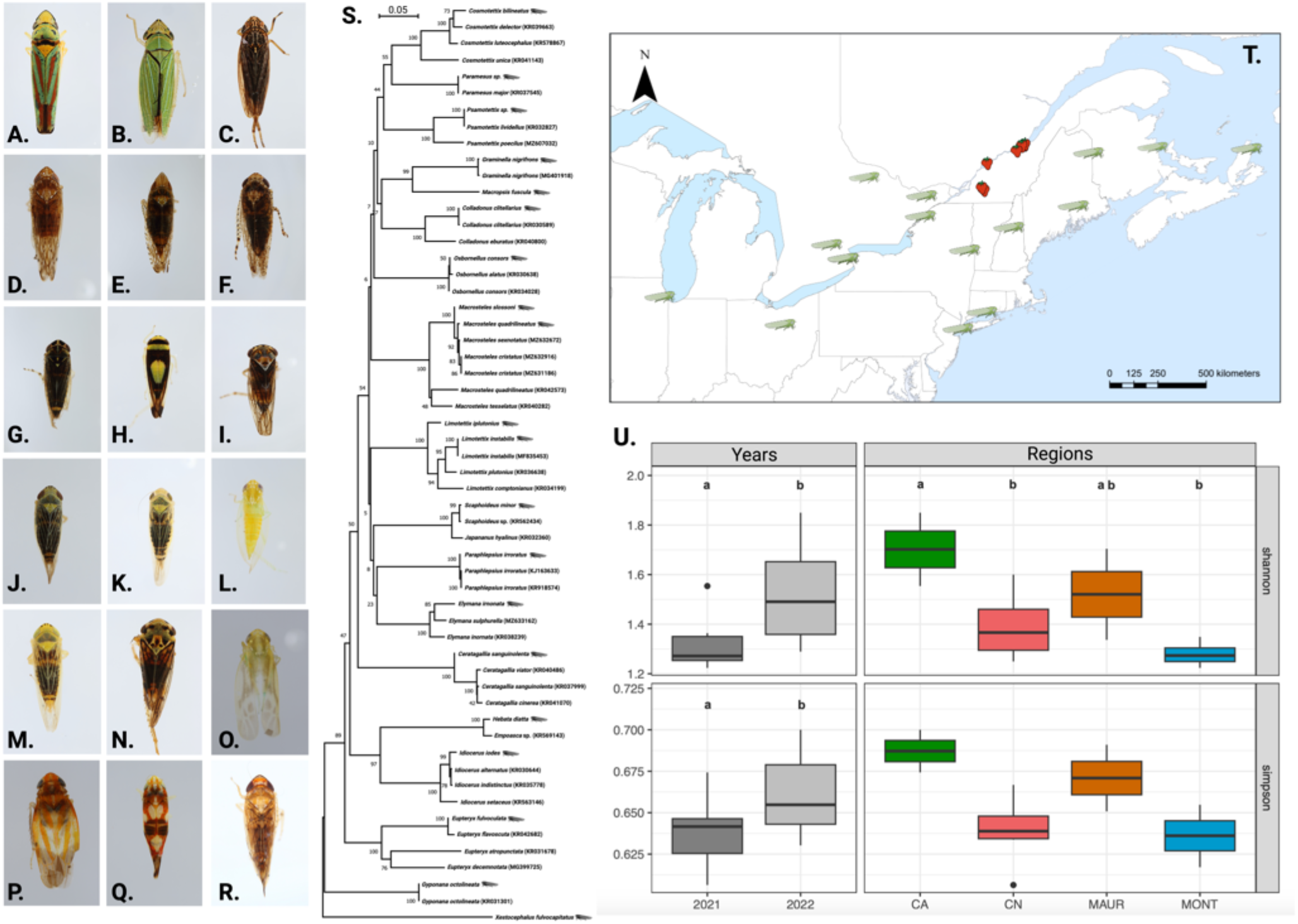
Leafhopper diversity reported in this study. **A**-**R** example of leafhoppers captured in strawberry fields with **A**, *Graphocephala coccinea*; **B**, *Draeculacephala noveboracensis*; **C**, Neokolla sp. ; **D**, *Osbornellus consors*; **E**, *Scaphytopius acutus*; **F**, *Paraphlepsius* sp. ; **G**, *Macrosteles* sp. ; **H**, *Colladonus clitellarius*; **I**, *Idiocerus formosus*; **J**, *Graminella nigrifrons*; **K**, *Dikraneura mali*; L, *Empoasca fabae*; **M**, *Graminella palludila*; **N**, *Macropsis fuscula*; **O**, *Hebata vergena*; **P**, *Erythridula plena*; **Q**, *Erythroneura elegans*; **R**, *Osbornellus limosus*. **S**, phylogenetic analysis of CO1 sequences generated from 20 of the 168 leafhopper species captured during this study (grey insect). GenBank accession number is provided between parenthesis for those sequences retrieved from the database. Bars represent 5 nucleotide substitutions in 100 positions. **T**, the latest report for the new leafhoppers reported for eastern Canada and Canada identified by leafhoppers in relationship with the strawberry fields screened in this study identified by strawberries. This localization used data from the Canadian National Collection of Insects, Arachnids and Nematodes and records available through different taxonomic key databases such as 3I (http://dmitriev.speciesfile.org/3i_keys.asp). **U**, alpha diversity Shannon and Simpson comparison between growing seasons and among regions. Letters indicate significant differences between years with *p* = 0.0151 for Shannon and *p* = 0.0417 for Simpson. Significant differences were only detected among regions for Shannon, with *p* = 0.0275, but not for Simpson, with *p* = 0.1614. Regions are differentiated by colour and code with green and CN for Capitale-National, red and CA for Chaudière-Appalache, orange and MAUR for Mauricie, and blue and MONT for Montérégie.

### 3.2. Population distribution and influence of temperature

The weekly mean temperatures recorded for the 247 days of leafhopper capture ranged from 6.90 °C to 24.6 °C, with a median of 18.5 °C. The chosen model for all leafhopper species abundance used a positive linear function with the degree-days base 10 °C, meaning that an increase in daily mean temperatures within the range of temperatures registered during this experiment caused an increase in leafhopper abundance (Fig. 3A). However, for *E. fabae* and *M. quadrilineatus,* the best models used the mean temperature as their temperature-related index because it was more parsimonious (Fig. 3A-B). For the overall leafhopper population, *E. fabae* abundance also followed a positive linear function with temperature (Fig. 3C). However, *M. quadrilineatus*’ relation to temperature was best modelled with a quadratic equation, and the abundance increased until 21°C but then decreased (Fig. 3B, Supplementary Table S11). When the effect of temperature was removed, the overall leafhopper abundance increased until the second half of July and then decreased, following a quadratic function (Fig. 3D). The distribution of *M. quadrilineatus* followed a positive linear function after removing the effect of temperature (Fig. 3E), while the distribution of *E. fabae* followed a cubic equation, and the peak abundance was observed earlier at the beginning of July, with another increase observed by the second half of August (Fig. 3F).

**Figure 3.**
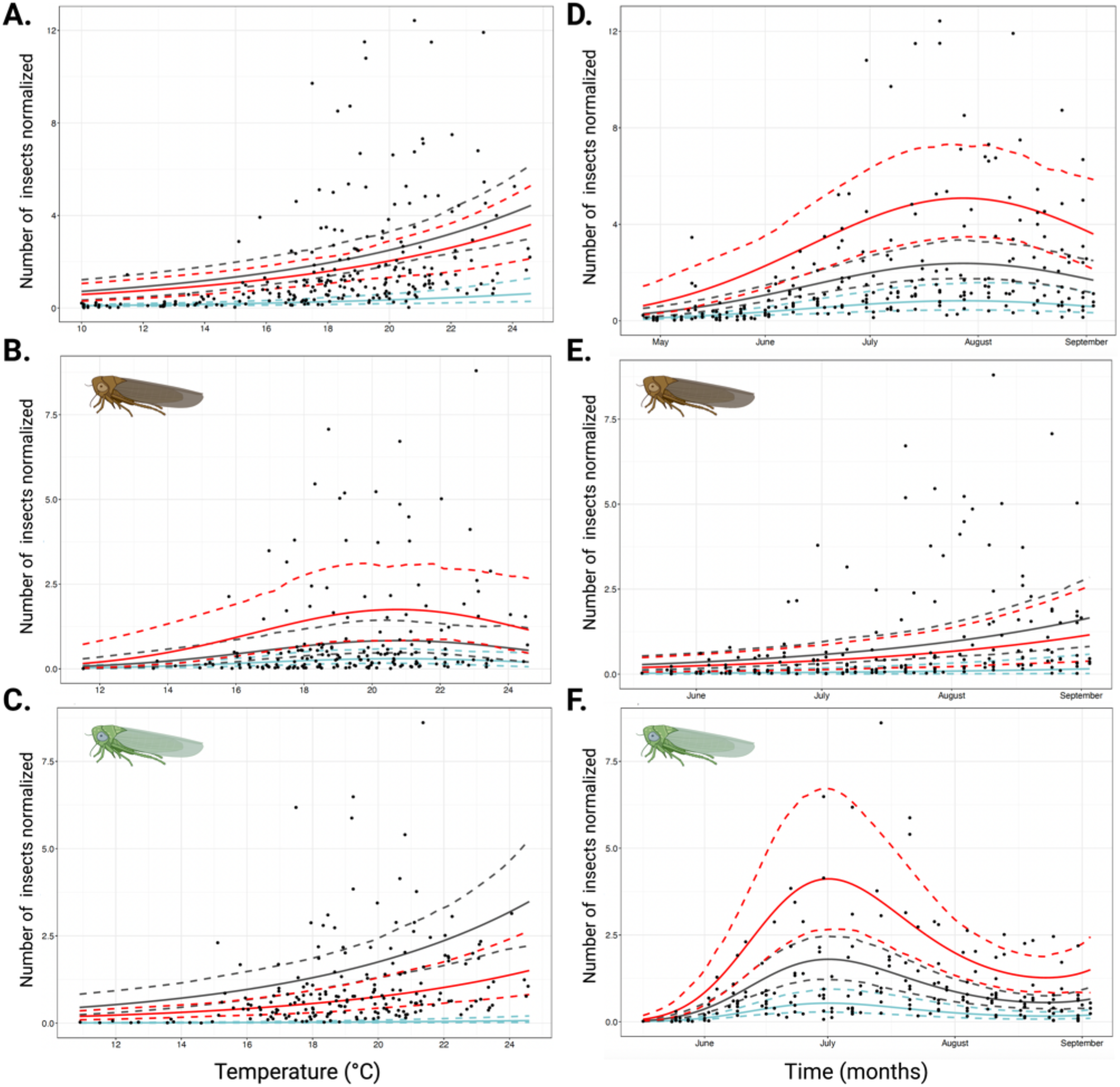
Leafhopper population structure. **A**-**C** influence of temperature on **A**, global leafhopper population; **B**, *Macrosteles quadrilineatus*; and **C**, *Empoasca fabae*. **D**-**F**, Structure of the population after removing the effect of temperature on **D**, global leafhopper population; **E**, *M. quadrilineatus*; and **F**, *E. fabae*. The model for the global leafhopper population was α(*) + β1·day (***) – β2·day2 (***) + β3·base10 (***); α + β1·day (***) + β2·T°C2 (*) + β3·T°C (**) for *M. quadrilineatus*; and - α + β1·day - β2·day2 (***) + β2·day3 (***) + β3·T°C (***) for *E. fabae*, where * p-value ≤ 0.05, ** p-value ≤ 0.01, and *** p-value ≤ 0.001. Day = number of days since January 1^st^ and T°C = mean daily temperature (°C). The red line represents the maximum value of days or temperature in the data used for the modelling, the black line is the median, and the blue line is the minimum value (Supplementary Table S11).

Models for overall leafhopper abundance and for *E. fabae* were highly significant, with *p*-values generally lower than 1x10^-5^. For *M. quadrilineatus,* the model was less significant, with p-values lower than 1x10^-6^ for the time factor and higher p-values for the temperature factor (*p* < 0.024). Weekly precipitation did not contribute to the model, and no relevant collinearity was found between the rainfall and the mean temperature, with Pearson’s coefficient = 0.088.

### 3.3. SbGP phytoplasma and disease incidence

Typical symptomatology associated with SbGP disease was observed during both sampling seasons (Fig. 4A-B). These constitute two of the sixteen SbGP reports registered in Québec in the last ten years distributed around the province, with a clear increase in the last three years (Fig. 4C). A 1.25 kb band corresponding to the F2nR2 fragment was observed through 1% agarose gel electrophoresis when using DNA extracts from symptomatic plants, while no amplification was obtained from asymptomatic plants. The F2nR2 sequences obtained shared more than 99% similarity with the reference strain for the 16SrI group, the Aster Yellows (AY, GenBank accession no. M30790), reference strain for the species ‘*Candidatus* Phytoplasma asteris’, through BLAST. This result was supported by *in silico* RLFP analysis using *i*PhyClassifier, which identified the strains as members of group 16SrI (Fig. 4D). Two distinct F2nR2 sequences were retrieved from the samples. One of the sequences was identified as 16SrI-S by the RFLP digestion pattern (Fig. 4D), and the coefficients of similarity for these sequences were 1.00 with the reference strain LiLL (16SrI-S, GenBank accession number HM067755). The other F2nR2 sequences were a variant of 16SrI-R showing a coefficient of similarity of 0.98 with the reference strain ChBL (16SrI-R, GenBank accession number HM067754) (Fig. 4D). We also determined that among those sequences identified as 16SrI-S and 16SrI-R, only the restriction pattern obtained with *AluI* is different, and it is responsible for the different subgroup classifications (Fig. 4E). Through *in vitro* restriction digestion of SbG-clone2 and SbG-clone5 amplicons with *AluI*, we were able to replicate the *in silico* pattern, confirming the results (Fig. 4E). The results obtained using *i*PhyClassifier were confirmed by phylogenetic analysis (Fig. 4F). The sequences classified as 16SrI-S grouped with the reference strain LiLL, while the sequences classified as the 16SrI-R subgroup grouped with the reference strain ChBL (Fig. 4F).

**Figure 4.**
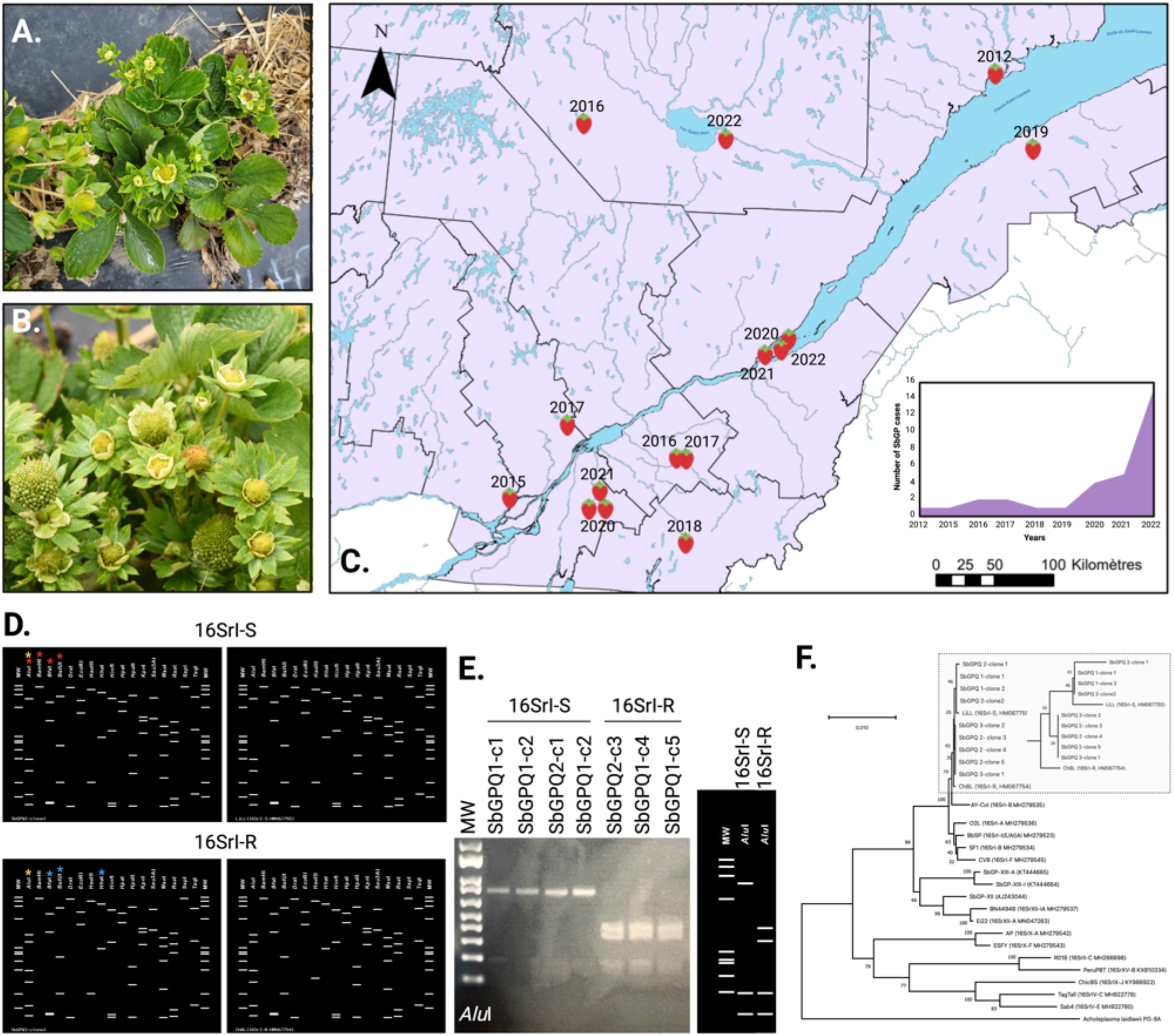
Distribution and characterization of strawberry green petal disease in Québec. **A**-**B**, Typical strawberry green petal disease (SbGP) symptoms observed in Québec. Green flowers and the presence of a green fruit-like structure. **C**, map with the distribution of all the SbGP cases in the province of Québec in the last ten years. The geographic location of the cases is represented by a strawberry, and the report’s year is also presented on the map. The graph placed in the right bottom corner represents SbGP incidence from 2012 to 2022. **D-F**, characterization of the SbGP strain affecting strawberry plants in Québec. **D**-**E**, distinctive RFLP patterns for the new 16SrI-(S/R)S subgroup delineated in this study. **D**, RFLP patterns obtained with *i*PhyClassifier from *in silico* digestion of 16S rDNA gene F2nR2 fragments from SbGPQ clones, strains LiLL (16SrI-S), ChBL (16SrI-R), and AY-Col with a set of 17 enzymes previously defined in the scheme of phytoplasma classification. Red stars mark differences among 16SrI-S strains and AY-Col, blue stars mark differences among 16SrI-R strains and AY-Col, and yellow stars mark differences among 16SrI-S and 16SrI-R strains. **E**, *In vitro* confirmation of the differential RFLP obtained after F2nR2 digestion with *Alu*I among 16SrI-S SbG clones and 16SrI-R SbG clones. MW represents the 1 kb plus pattern. **F**, phylogenetic analysis of the 16S operon obtained from different clones sequenced from samples SbG-3, branching with phytoplasma strains into the 16SrI-S and 16SrI-R subgroups. The bar represents 1 nucleotide substitution in 100 positions, and the square also encloses the branch grouping clones of the SbGPQ strain.

The 600 bp *cpn60* UT sequence was amplified from all symptomatic samples. The *cpn60* UT sequences generated in this study shared 100% sequence identity among them, and BLAST analysis showed that SbGPQ *cpn60* UT sequences shared 100% identity with the AY-Col strain (16SrI-C, GenBank accession no. KJ939994). This result was consistent with the *Cpn*ClassiPhyR classification, suggesting that the SbGPQ phytoplasma strain is a member of the cpn60UT I-IC subgroup, sharing the same RFLP digestion pattern (Supplementary Fig. S2). This classification was supported by phylogenetic analysis using the sequences obtained from each sample and other sequences from 16SrI subgroups. Finally, the heterogeneity of the SbGPQ phytoplasma strain was confirmed by hybridization-based multilocus sequence typing, finding that three of the samples analyzed had a 16SrI-R operon, while eight had a 16SrI-S operon (Supplementary Fig. S3 and Table S12). However, the rest of the markers did not show the same level of heterogeneity observed in the 16S operon, grouping with 16SrI phytoplasma strains (Supplementary Fig. S3). Following the nomenclature suggested for heterogeneous phytoplasma groups, the phytoplasma strain detected is a member of the heterogeneous novel 16SrI-(S/R)S phytoplasma subgroup.

### 3.5. Leafhopper microbiome

No phytoplasma was detected in the 3 824 leafhoppers analyzed in this study, including those captured with sticky yellow traps or with the aspirator. Similarly, although the plants incubated with the leafhoppers showed clear evidence of feeding (Supplementary Fig. S4), no phytoplasma was detected in the plant tissue. These results were consistent with the analysis of *M. quadrilineatus*, the aster leafhopper, microbiome, where we did not find any OTU corresponding to phytoplasmas in any of the 60 samples analyzed, which grouped a total of 600 specimens (Fig. 5A-C).

**Figure 5.**
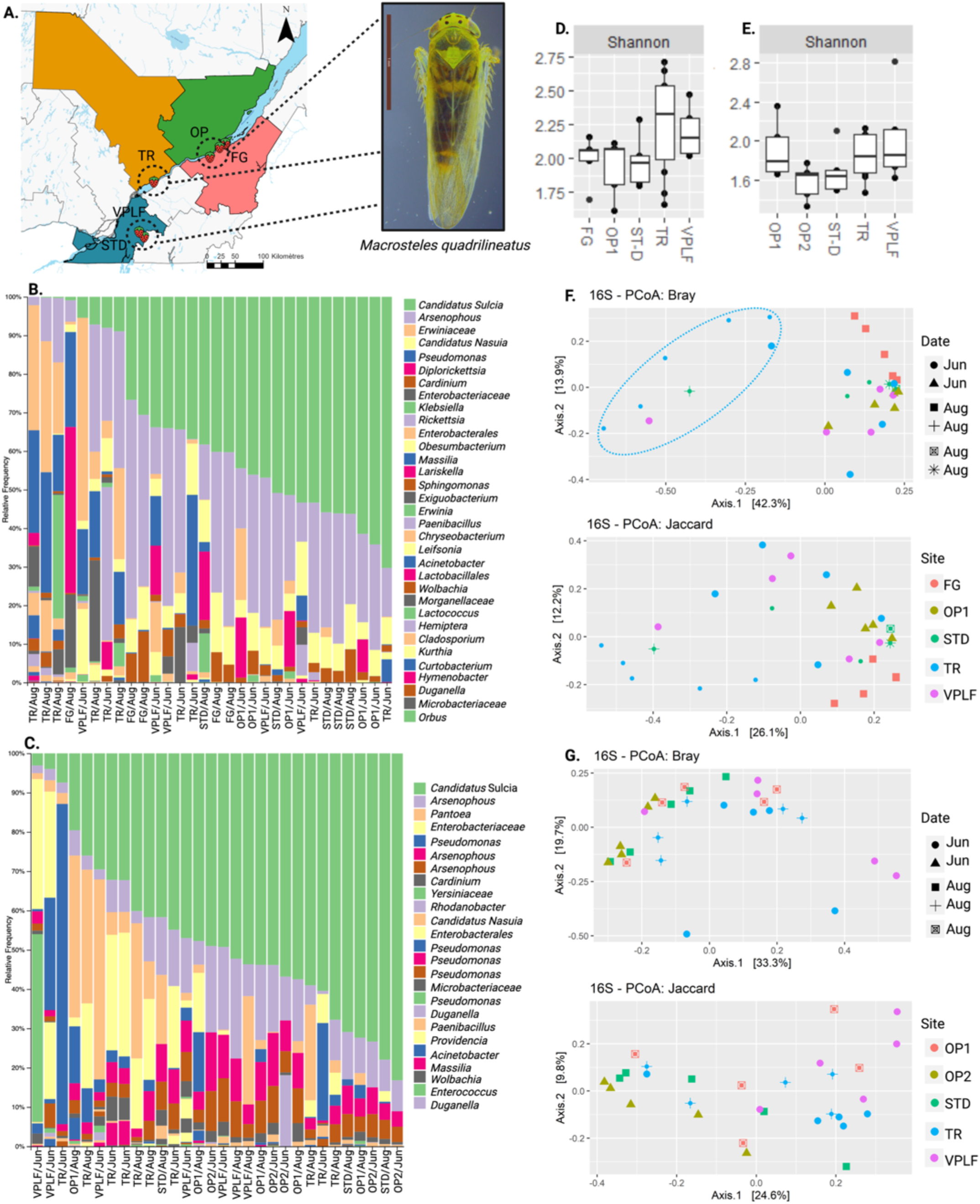
*Macrosteles quadrilineatus* microbiome analysis. **A**, location of the sites where the specimens tested were captured. OP, FG, TR, VPLF, and STD are codes used to identify strawberry farms located in three different geographic regions, as presented in figure 1. **B**-**C**, Relative abundance of operational taxonomic units at the genus level for both growing seasons with **B** for 2021 and **C** for 2022. **D**-**E**, alpha diversity comparison among microbial communities in insects sampled at different farms with **D** for 2021 and **E** for 2022. No statistical significance was found with *p* < 0.05. **F**-**G,** Distribution of the samples according to beta diversity calculated with Bray–Curtis and Jaccard indexes for 2021 and 2022, respectively. The forms represent the month of capture and the colour fields where the leafhoppers were captured.

The sequencing of the V4-V5 region of the 16S rRNA universal target on the aster leafhopper produced, after filtering out insect sequences and rare OTUs, a total of 344 992 sequences belonging to 153 different OTUs for those insects collected during the first growing season and a total of 212,630 sequences belonging to 129 OTUs for those insects collected during the second growing season (Fig. 5B-C, Supplementary Table S13). Globally, the diversity of the aster leafhopper microbiome between both growing seasons was very similar, a trend that was also observed among the geographical locations where the leafhoppers were collected (Fig. 5B-C). This was consistent with the observed, Shannon, and InvSimpson alpha diversity indices, which did not show statistically significant differences among the sites, indicating that the geographical gradient used in this study does not influence the alpha diversity of the bacterial community in the aster leafhopper (Fig. 5D-E, Supplementary Fig. S5). All detected OTUs were assigned to one of the four phyla Proteobacteria, Bacteroidota, Firmicutes, and Actinobacteriota, represented by eleven orders: *Pseudomonadales*, *Cytophagales*, *Flavobacteriales*, *Enterobacterales*, *Diplorickettsiales*, *Burkholderiales*, *Rickettsiales*, *Exiguobacterales*, *Paenibacillales*, *Micrococcales*, and *Orbales* (Fig. 5B-C, Supplementary Table S13). In each growing season, the most abundant sequences belonged to the obligate symbionts ‘*Candidatus* Sulcia’ spp. with 129,133 sequences and 27 OTUs in the first growing season and 105848 sequences and 56 OTUs in the second growing season; and ‘*Candidatus* Nasuia’ spp. with 23 567 sequences and 11 OTUs in the first year, but this number was surprisingly lower in the second year with 1, 720 sequences and only two OTUs (Supplementary Table S14). Two other genera that were highly represented in the aster leafhopper microbiome were *Arsenophonus* spp. and *Pseudomonas* spp., with 126 188 sequences and 28 OTUs and 39,030 and 35 OTUs, respectively (Supplementary Table S14). However, the core microbiome for the aster leafhopper was consistent between both growing seasons, formed by the genera ‘*Candidatus* Sulcia’, ‘*Candidatus* Nasuia’, the endosymbiont *Arsenophonus* spp. and the parasitic ‘*Candidatus* Cardinium’ (Supplementary Table S14).

To compare the microbial community structure among the aster leafhoppers collected in June and August and those collected in each sampling region, principal coordinate analysis (PCoA) of beta diversity analysis was performed based on Bray-Curtis dissimilarity and Jaccard similarity for each growing season (Fig. 5F-G). The analysis showed that neither the time of year nor the geographical location were major drivers of diversity among the microbial communities (Fig. 5F-G). However, based on the Bray-Curtis index for the aster leafhoppers captured during the first growing season, we can see some differentiation of the bacterial communities in those leafhoppers collected further south in June from the rest; however, this result was not significantly different (Fig. 5F).

### 3.6. Ineffectiveness of insecticides to control leafhoppers

After interviewing strawberry growers, we discovered that during both growing seasons, at least 12 different insecticides certified as efficient against leafhoppers were used (Supplementary Table S8). Of those, four were used more than five times and were used for statistical analysis comparing the leafhopper abundance before and after the insecticide treatment (Fig. 6A-B). The products selected to test their efficiency were Cormoran (acetamiprid and novaluron) with 10 applications, Assail (acetamiprid) with nine applications, Up-Cyde and Mako (cypermethrin) with five applications, and Malathion (organophosphorus) with five applications. We found no significant differences for any of the treatments when comparing before and after the application of the insecticide (*p* = 0.8488) (Fig. 6C, Supplementary Fig. S6). The same analysis was performed with the two most abundant leafhoppers, *E. fabae* and *M. quadrilineatus*, and the same results were obtained, confirming no reduction in the abundance of these species after insecticide application, with *p* = 0.6540 for *E. fabae* and *p* = 0.1781 for *M. quadrilineatus* (Supplementary Fig. S7).

**Figure 6.**
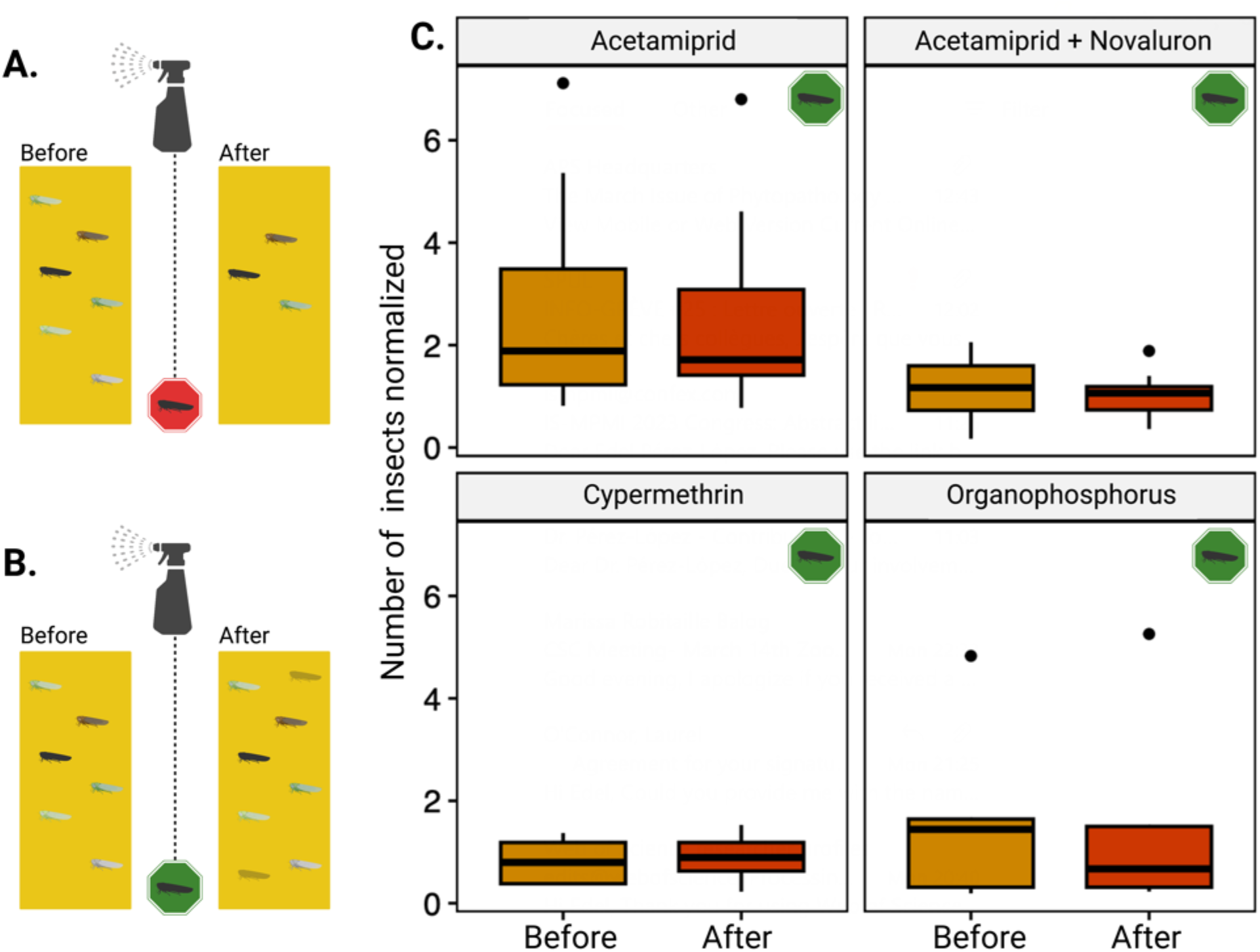
Effect of insecticides on leafhopper population control. **A**-**B** represent the analysis performed to identify the effectiveness of the insecticides used by strawberry growers. If the insecticide can reduce leafhopper abundance, it would receive a red tag; if not, it would receive a green tag. **C**, none of the insecticides applied by the growers and analyzed statistically (n ≥ 5 applications) had a significant reduction in the abundance of leafhoppers (*p* = 0.8488).

## 4. DISCUSSION

The effect of climate change on agriculture is complex and multilayered (Wagner et al., 2021; Jägermeyr et al., 2021). The effects of different climate change-related stressors on agricultural pests and insect-transmitted diseases have been poorly studied, and we lack comprehensive models to advance our understanding. This study provides indications that Nearctic leafhoppers can be used as a sentinel to investigate those phenomena, considering the rapid rise in temperatures experienced in this geographic region and their ability to transmit devastating plant diseases (Islam et al., 2020; Brochu et al., 2023).

Consistent with what the growers reported to us during initial interviews and with previous studies performed on small fruits in eastern Canada, the first leafhopper peak of abundance was observed in June (Pagé, 2013; Saguez et al., 2014). Later, two other peaks of abundance occurred in July and August, corresponding to the higher temperatures registered in the seasons, which could be attributed to the arrival of migratory species such as the potato leafhopper *Empoasca fabae* and the aster leafhopper *Macrosteles quadrilineatus*. Overall, 118 leafhopper species were identified in strawberry fields, which is 102 more species than previously reported 14 years before in vineyards (Saguez et al., 2014) and 53 more species than previously reported 10 years before in blueberry fields (Pagé, 2013) in Québec. Although these studies are hard to compare considering the sampling methods, site selection, hosts, and exhaustive identification of the specimens collected in this study, we found that based on the Shannon diversity index, leafhopper communities in vineyards were more diverse in Québec 14 years ago than the communities found in strawberry fields in 2021 and 2022 (Supplementary Fig. S8).

Surprisingly, most of the new reports for eastern Canada found in strawberry fields are members of the genera *Erythridula* and *Erythroneura,* key pests of vineyards, and not migratory species that were previously reported in the northern United States and in Ontario but never as far east as Québec (Dmitriev & Dietrich, 2009; Jarrell et al., 2020). With increasing temperatures and climate change events, some leafhopper species may have been able to expand their ecological niche and their distribution areas (Santana et al., 2019). Our results demonstrate that several nonmigratory Nearctic species are settling in further northern regions, probably by expanding their thermal limits. These rare species have a strong influence on the Shannon index and could be what induced differences in the diversity among the geographic regions studied, in addition to different agricultural practices performed by each grower, a factor that reshapes insect diversity (Outhwaite et al., 2022). Previous studies have shown that host plants in the landscape have a strong influence on the biodiversity and the number of leafhopper species present at a site (Helbing et al., 2017; Dainese et al., 2015; Tscharntke et al., 2012). The intensification of agriculture and cultivated fields around the sites can also affect the biodiversity observed by reducing the presence of rich natural habitats (Outhwaite et al., 2022).

In both growing seasons, *E. fabae* and *M. quadrilineatus* represented 88.4% of the captures during the first growing season and 77.4% during the second. These two leafhopper species are well-known phytoplasma vectors and are classic examples of migratory pests in the Nearctic region, extending from the southern United States to eastern Canada (Baker et al., 2015; Capinera, 2008). Unlike most Nearctic leafhoppers, these two species are multivoltine, with multiple generations per year, strongly depending on climatic conditions (Hogg and Hoffman, 1989; Capinera, 2008; Chasen et al., 2014). Both species were first captured during the last week of May in both growing seasons, from 11 to 22 days before the *E. fabae* average arrival dates reported for the states of New York and Massachusetts from 1951 to 2012 (Baker et al., 2015). Although there are no studies on the influence of temperature on aster leafhopper migration, the potato leafhopper migratory pattern is strongly influenced by temperature, precipitation and agricultural practices, and higher temperatures influence earlier arrival and severity of leafhopper infestation (Maredia et al., 1998; Baker et al., 2015; Chen et al., 2019).

Although moisture stress has been identified as playing a role in potato leafhopper severity in crops such as alfalfa (Schroeder et al., 1988) and precipitation has a major influence on insect development (Daane & Williams, 2003; Chen et al., 2019), the models did not show a clear relationship with the mean total weekly precipitation. Curiously, during the second growing season, the year with higher precipitation, the global abundance of the potato leafhopper was 10% lower. The index used for *M. quadrilineatus* and *E. fabae* did not significantly influence the modelling because they are migratory species that arrive when temperatures are over 10 °C. In this study, they were first detected when the weekly mean temperature was 10.9°C. However, the choice of a temperature-related index may be more relevant for migratory species when modelling past September when the temperature starts decreasing. For all leafhoppers, the use of a degree-day base of 10 °C suggests that temperatures colder than 10 °C may slow nonmigratory leafhopper species reproduction or activity (Trudgill et al., 2005). This finding is supported by a previous study that found that leafhoppers commonly collected in vineyards in southern Québec are best modelled with a degree-days base 8 °C index (Bostanian et al., 2006).

With an increase in mean temperatures in Québec, the model for the global leafhopper population and for the potato leafhoppers suggests that leafhopper abundance will increase. In addition, these species do not encounter their optimal, or maximal, temperature in Québec, resulting in an expectation of an increase in their abundance. Unfortunately, there are no data available regarding *E. fabae* thermal limits, but as mentioned before, high temperatures favour this pest migration and development, confirming our models (Maredia et al., 1998; Baker et al., 2015; Chen et al., 2019). On the other hand, an increase in temperature had a mixed effect on *M. quadrilineatus*, with an expected increase in abundance early and late in the growing season but a decrease when the temperature was over 21 °C. Based on our model, an increase in the temperature will be disadvantageous for the aster leafhopper species, supported by the experimental results where the median temperature registered within our sampling sites was under the 20 °C threshold. Our model is also supported by a previous study that found that the aster leafhopper, under controlled conditions and in a preferential host such as barley, has the highest survival up to 20 °C, although some individuals could survive up to 30 °C, temperatures that seriously affect the reproduction of *M. quadrilineatus* (Bahar et al., 2018).

When the effect of the temperature is removed, the global leafhopper abundance peaks in mid-July, and for *E. fabae,* the peak is observed at the beginning of July. This may be caused by the overlap of migratory waves and native populations from different leafhopper species. The models also show that the decrease in abundance afterwards is caused not only by the decrease in the temperatures but also by other factors that were not included in the model. These fluctuations are not seen with *M. quadrilineatus*, whose population would continually increase without the effect of temperature. The model also suggests that the potato leafhopper population starts increasing again during the second half of August, but a longer period of sampling would be needed to investigate and confirm this tendency.

These two leafhopper species show different population structures over time and under the influence of temperature. Since they are dominant and major pests in this region, field management strategies should consider these species separately to achieve successful control (Zhou et al., 2003; Olivier et al., 2009; Pagé, 2013; Saguez et al., 2014; Chasen et al., 2014; Gavloski, 2018).

Despite our efforts, we were unable to identify the leafhoppers or leafhoppers transmitting SbGP phytoplasma in eastern Canada after analyzing 10% of the leafhoppers captured during the first growing season. The detection rate of phytoplasma in leafhoppers is low, with reports as low as 1.8% of samples positive after analyzing 11 515 leafhoppers captured in Canadian vineyards (Olivier et al., 2014). The low incidence of SbGP disease in the last decade explains why it might have been impossible to find positive leafhoppers, although it is clear that the number of cases has been growing from five to ten more times in the last couple of years. Strawberry green petals were one of the first phytoplasma-associated diseases reported worldwide (Baker, 1956) and the first one reported in Canada in 1955 (Chiykowsky, 1962), specifically in eastern provinces, including Québec. The disease was initially thought to be caused by a virus, but it has since been determined that SbGP is associated with infection by a phytoplasma (Lee et al., 2004). It was not until last year that the strain present in eastern Canada was partially characterized (Plante et al., 2022), and in this study, the SbGPQ phytoplasma strain was identified as a new heterogenous phytoplasma subgroup. Our results point to a fast evolution of the pathogen associated with SbGP. In North America, strains within the 16SrI-C and 16SrI-R subgroups have been linked to SbGP (Gundersen & Lee, 1996; Harrison et al., 1997; Jomantiene et al., 2002), but now we have identified a new subgroup phylogenetically close to those subgroups but clearly different.

Aster leafhoppers feed on and transmit aster yellow phytoplasma strains to approximately 300 plant species in 200 different plant families (Olivier et al., 2009; Stillson & Szendrei, 2020; Romero et al., 2023). In Canada, the aster leafhopper has been confirmed as the vector of aster yellow phytoplasma in canola fields in prairies and has been identified as positive for phytoplasma in vineyards and blueberry fields (Olivier et al., 2009, Pagé, 2013; Olivier et al., 2014). This is what motivated us to investigate the aster leafhopper microbiome to further try to find the presence of SbGP phytoplasma. Once again, we were not able to detect the presence of phytoplasma in *M. quadrilineatus*; however, we found several facultative symbionts previously reported in the aster leafhopper that have been previously identified as strawberry pathogens (Streten et al., 2005; Bull et al., 2009; Bressan et al., 2012). For example, Rickettsia, an endocellular alphaproteobacteria that has been previously reported in the *Macrosteles striifrons*, *Macrosteles sexanotatus*, and *M. quadrilineatus* microbiome (Ishii et al., 2013; Mao et al., 2018), has been identified as a strawberry pathogen associated with strawberry lethal yellows disease in Australia (Streten et al., 2005). A new *M. quadrilineatus* facultative symbiont reported here is Arsenophonus, a very diverse arthropod endosymbiont previously reported in *Macrosteles laevis* parasitizing the obligate endosymbiont ‘*Candidatus* Sulcia’ (Kobiałka et al., 2015). Arsenophonus, specifically ‘*Candidatus* Arsenophonus phytopathogenicus’, has been identified as a strawberry pathogen causing a disease resembling strawberry redness in Europe and Australia but never in North America (Bressan et al., 2012; Bressan, 2014). It is especially intriguing that this endosymbiont has not been found in *M. quadrilineatus* populations reared in other plant hosts, but here, when collected from strawberry fields, it was observed.

Planthopper microbiomes have also been recently linked to insecticide resistance and adaptation to climate change (Zimmer et al., 2018; Pang et al., 2018; Li et al., 2020; Zhang et al., 2021; Gupta et al., 2022; Zhang et al., 2023), but this has not been studied in leafhoppers. In the brown planthopper *Nilaparvata lugens*, for example, it has been found that the presence of two distinctive *Arsenophonus* strains can change the expression of cytochrome P450, CYP6ER1, which is responsible for imidacloprid detoxification and decreases insecticide resistance (Pang et al., 2018). *Arsenophonus*, a member of the aster leafhoppers captured in our study core microbiome, has been identified as a key member of the “insecticide-resistant microbiome” along with *Wolbachia*, *Acinetobacter rhizosphaerae*, and *Staphylococcus sciuri*, and the fungi *Hirsutella* spp. (Zhang et al., 2023). However, the host genotype background and relationship with the environment are factors that modulate this detoxifying microbiome-mediated effect (Li et al., 2020; Zhang et al., 2021; Zhang et al., 2023). Intriguingly, although it is difficult to draw conclusions on this subject from our microbiome data, we found that the insecticide treatments used by strawberry growers did not control leafhopper populations during the growing seasons analyzed. This tolerance or resistance to insecticide observed could also be a direct consequence of climate warming, promoting migrations or expansion of the overwinter range of pests, as observed for the diamondback moth *Plutella xylostella* (Ma et al., 2021), although there is no record of this phenomenon for leafhoppers. Something that should also be further investigated is the effectiveness of insecticides at early leafhopper developmental stages. Leafhopper nymphs are not mobile and are more likely to be effectively controlled than adults (O’Hearn et al., 2017; Tirello et al., 2021).

## 5. CONCLUSIONS

The use of insects as sentinels of global changes has been previously suggested; however, their high diversity, rapid population dynamics and the paucity of information regarding several taxa have also been identified as challenges (Wilson and Fox, 2020). Our results suggest that migratory Nearctic leafhoppers such as the aster leafhopper and the potato leafhopper can be used as a model to study the effect of climate change on (*i*) insect migratory patterns, (*ii*) insect population dynamics, (*iii*) biological invasions, (*iv*) disease dispersion, (*v*) pathogen evolution, and (*vi*) insecticide resistance. We recommend further studying this group of insect pests to gain a better understanding of their ecology and mechanism affected by environmental stressors and fully exploit their potential as possible climate change indicators. If achieving sustainable agriculture is a challenge, achieving sustainable agriculture in a changing climate is a colossal task that requires all the tools at our disposal.

## ACKNOWLEDGEMENTS

This work was funded by MAPAQ through the Program Innov’Action Agroalimentaire, project number IA120641. Thank you to Centre Sève, FRQNT for supporting N.P. internship at the University of Florida to receive leafhopper taxonomy training, to NSERC to support J.D. summer internship in our lab through the USRA program and to Mitacs for supporting D.T. through the Globalink program. Thank you to the RQRAD, MAPAQ and FRQNT for supporting the publication of this work as open access. The authors would like to thank Gaétan Daigle for the statistics assessment and all the strawberry growers and agronomists who have collaborated and continue to collaborate with our group.

## AUTHOR CONTRIBUTIONS

Conceptualization: C.G., S.T., V.F., E.P.-L. Data curation: N.P., J.D., A.-S.B., T.D., A.D., E.P.-L. Fieldwork: N.P., F.M., D.T. Insect identification: N.P., J. K., B.B., J.-P.L. Laboratory analyses: N.P., J.D., A.-S.B., T.D., D.T. Visualization, statistical analysis, and modelling: J.D., T.D., E.P.-L. Project administration: C.G., V.F., E.P.-L. Resources: C.G., V.F., S.T., A.D., E.P.-L. Supervision: V.F., E.P.-L. Writing: N.P., J.D., T.D., E.P.-L.

## CONFLICT OF INTEREST

The authors declare that they have no conflicts of interest.

## DATA AVAILABILITY STATEMENT

The code used in this study for statistical analysis and modelling can be found here https://github.com/Edelab/Leafhoppers/tree/main. The code used for the microbiome analysis can be found here https://github.com/kevmu/dada2_analysis_pipeline. The phytoplasma-derived sequences were deposited in GenBank under the accession numbers ON511092 to ON511100 and OR074733 to OR074743 for phytoplasma 16Sr operon; ON376316 to ON376321 for phytoplasma *cpn60* UT, OR078459 to OR078469 for phytoplasma *secY*; and OR078470 to OR078481 for phytoplasma *tuf*. The microbiome data sequence files generated during the current study are available in the National Center for Biotechnology Information Short Read Archive under BioProject PRJNA977644 (https://www.ncbi.nlm.nih.gov/sra/PRJNA977644). Supplementary Tables can be found here https://doi.org/10.5281/zenodo.8025215. Supplementary Figures can be found here https://doi.org/10.5281/zenodo.8025512. The summary and conclusions of the study in French can be found here https://doi.org/10.5281/zenodo.8025729.

## ORCID

*Tim Dumonceaux* https://orcid.org/0000-0001-5165-0343

*Brian Bahder* https://orcid.org/0000-0002-1118-4832

*Joel Kits* https://orcid.org/0000-0003-2685-0567

*Antoine Dionne* https://orcid.org/0000-0003-4359-2467

*Stéphanie Tellier* https://orcid.org/0000-0002-4191-2706

*Charles Goulet* https://orcid.org/0000-0002-3833-1492

*Valérie Fournier* https://orcid.org/0000-0003-0711-5241

*Edel Pérez-López* https://orcid.org/0000-0002-3708-8558

## Notes

https://www.ncbi.nlm.nih.gov/sra/PRJNA977644

https://doi.org/10.5281/zenodo.8025215

https://doi.org/10.5281/zenodo.8025512

https://doi.org/10.5281/zenodo.8025729

### Competing Interest Statement

The authors have declared no competing interest.

